# Normalization models of contextualized touch

**DOI:** 10.1101/283218

**Authors:** Md. Shoaibur Rahman, Jeffrey M. Yau

**Affiliations:** Department of Neuroscience, Baylor College of Medicine, Houston, TX 77030 USA

**Author notes:** Corresponding author: Jeffrey M. Yau One Baylor Plaza, T111 Houston, TX 77030.

## Abstract

Divisive normalization is a canonical computation that explains contextual modulation of visual perception and neural responses in the visual system. Conceivably, normalization also underlies contextual modulation of bimanual touch, a perceptual process that likely requires combining what is felt on the hands with where the hands are located in space. We found that touch experienced on one hand systematically modulates how touch is perceived on the other hand. Notably, bimanual interaction patterns and their sensitivity to hand locations differed depending on whether participants directed attention to the frequency or intensity of the cues, which were always mechanical vibrations. These idiosyncratic perceptual patterns were well explained by distinct cue combination models that each comprise divisive normalization. Our findings indicate that, while feature-specific rules govern bimanual touch, normalization underlies contextual modulation between the hands.

**Significance Statement:** How we perceive sensory cues depends on the context in which we experience them. Contextual modulation of vision results from divisive normalization, a canonical computation which adjusts the activity of visual neurons according to the pooled activity over the neural population. We tested the hypothesis that contextual interactions between cues felt on the two hands are also consistent with normalization. We found that touch on one hand systematically influenced perception on the other hand. Moreover, we observed distinct contextual modulation patterns when subjects attended to the frequency or intensity of the cues, which were always mechanical vibrations. Despite these differences, normalization models accounted for both perceptual patterns. Our results support the notion that normalization underlies contextual modulation between the hands.

## Introduction

How we perceive and understand sensory information depends on the context in which we experience the sensory cues. For instance, the visual perception of stimulus contrast, tilt, and motion depend strongly on contextual spatial factors defined by the properties of the visual surround (1–3). These instances of contextualized perception are thought to be a consequence of spatial context processing in neural populations encoding visual features: the activity of an individual neuron is modulated by the neuronal population activity in which that neuron is embedded. Such gain control, termed divisive normalization (4), is thought to be a canonical computation performed by neural circuits (5). Indeed, normalization has also been invoked to explain contextual processing in audition (6), olfaction (7), and even value-guided decision making (8,9). Surprisingly, there have been limited efforts to relate touch and somatosensory operations to normalization (though see (10)). Spatial contextual processing in touch represents a unique challenge for the nervous system because the somatosensory system contains a deformable sensory sheet – the relative locations of touch receptors can change depending on the positioning of the limbs and hands. Accordingly, contextual processing of tactile cues on the two hands likely requires combining what is felt on the hands with where the hands are located in space.

Here, we tested the hypothesis that contextual processing in touch is supported by divisive normalization. To probe contextual processing in touch, we characterized how subjects discriminate touch under bimanual stimulation and determined whether cutaneous interactions depend on the locations of the hands. We predicted that the experience of tactile cues on one hand would be contextualized by cues signaled on the other hand and the relative locations of the hands. We found that bimanual cue combination was obligatory regardless of whether subjects attended to vibration frequency or intensity; however, the specific contextual modulation patterns as well as their sensitivity to hand location differed depending on the attended feature. Critically, the idiosyncratic perceptual interactions were explained by distinct cue combination models that each involved divisive normalization.

## Results

To characterize bimanual contextual modulation in the frequency domain, we had participants perform a frequency discrimination task using their right hand as we manipulated the frequency of a distractor cue presented to their left hand and the location of the left hand. While participants maintained high performance levels in all conditions, the distractor cues systematically and reliably altered response patterns despite an explicit instruction to ignore the distractors (**Fig. 1*A*** and **Fig. S1**). The perceived frequency of the 200-Hz target stimulus (**Fig. 1*B***) was significantly biased toward distractor frequencies (frequency main effect: *F*_2,14_ = 147.0, *P* = 4.0e-10, η_p_^2^ = 0.95) in a manner that depended on distractor hand location (location effect: *F*_2,14_ = 6.9, *P* = 0.008, η_p_^2^ = 0.5; interaction effect: *F*_4,28_ = 17.8, *P* = 2.3e-7, η_p_^2^ = 0.72). Distractors also significantly altered perceptual thresholds (**Fig. 1*C***) in a manner that depended on frequency and hand location (frequency main effect: *F*_2,14_ = 12.6, *P* = 0.0007, η_p_^2^ = 0.64; location main effect: *F*_2,14_ = 0.6, *P* = 0.55, η_p_^2^ = 0.08; interaction effect: *F*_4,28_ = 12.0, *P* = 8.6e-6, η_p_^2^ = 0.63). When the target and distractor frequencies differed, distractors induced greater biases in frequency judgments and elevated discrimination thresholds more as the hands were located closer in space. When the target and distractor frequencies matched, target frequency estimates remained unbiased, but discrimination thresholds increased with larger separations between the hands. Importantly, merely manipulating the location of the left hand without delivering distractor cues did not alter right hand discrimination performance (**Fig. S2**). These patterns reveal that tactile frequency perception on right hand is selectively contextualized by what was experienced on the left hand and the left hand’s location.

**Figure 1.**
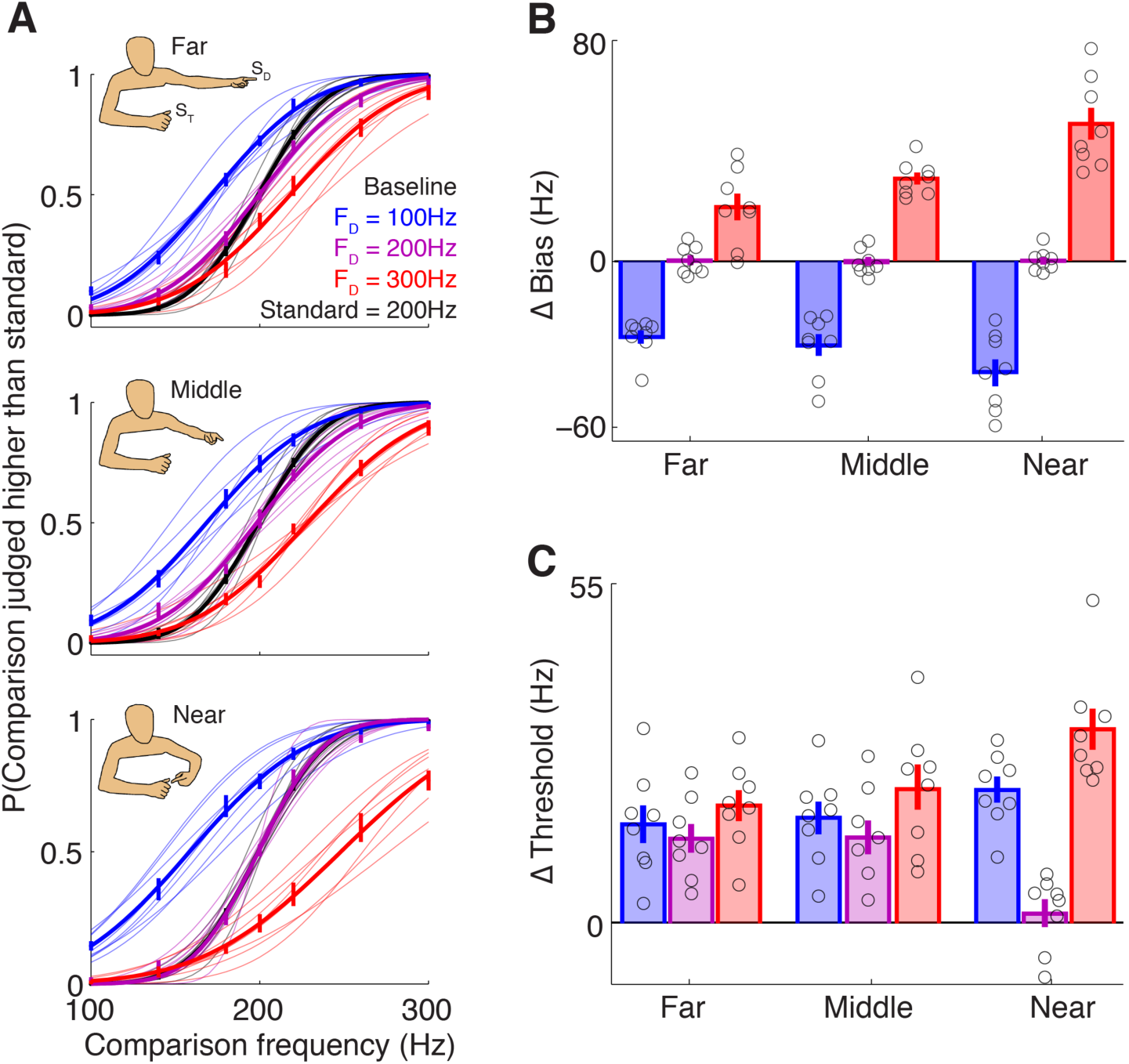
Bimanual interactions in frequency perception. (*n* = 8) (***A***) Average choice probability and psychometric curves from the group (thick traces) and individual subjects (thin traces). Target stimulus (S_T_) perception in the presence of distractor stimuli (S_D_) differed from perception without distractors (Baseline) depending on distractor frequency (f_D_) and distractor hand location. (***B***) Changes in bias (PSE), with respect to baseline, with each distractor frequency and location. Markers indicate individual subjects. Error bars indicate s.e.m. (***C***) Changes in discrimination threshold (JND) with respect to baseline.

To characterize bimanual contextual modulation in the intensity domain, participants performed an analogous intensity discrimination task in which distractor amplitude was manipulated in addition to distractor hand location. If left hand distractors similarly contextualize cue perception on the right hand for signals in the intensity and frequency domains, we predicted that 1) the perceived intensity of a target cue would be biased toward the intensity of the distractor cues, and 2) the magnitude of distractor effects would vary according to hand location. Although distractors systematically and reliably altered the perceived intensity of the target cue (**Fig. 2*A*** and **Fig. S1**), the pattern of distractor effects in the intensity discrimination task clearly differed from the pattern observed in the frequency discrimination task. Distractors only reduced the perceived intensity of target cues (**Fig. 2*B***) and the magnitude of attenuation scaled with distractor amplitude (amplitude main effect: *F*_2,14_ = 12.4, *P* = 0.0008, η_p_^2^ = 0.64). Notably, distractor effects did not vary with hand location (location main effect: *F*_2,14_ = 0.3, *P* = 0.74, η_p_^2^ = 0.04; interaction effect: *F*_4,28_ = 0.2, *P* = 0.94, η_p_^2^ = 0.03). Moreover, distractors exerted no consistent influences on intensity discrimination thresholds (**Fig. 2*C***) (amplitude main effect: *F*_2,14_ = 1.7, *P* = 0.23, η_p_^2^ = 0.19; location main effect: *F*_2,14_ = 0.04, *P* = 0.96, η_p_^2^ = 0.005; interaction effect: *F*_4,28_ = 2.1, *P* = 0.11, η_p_^2^ = 0.23). Thus, tactile intensity perception on the right hand is modulated by signals on the left hand, but in manner that is invariant to hand position changes. Because contextual modulation patterns in vibrotactile perception differ markedly depending on the attended feature, we infer that the nervous system employs distinct computations to support bimanual cue combination in the frequency and intensity domains.

**Figure 2.**
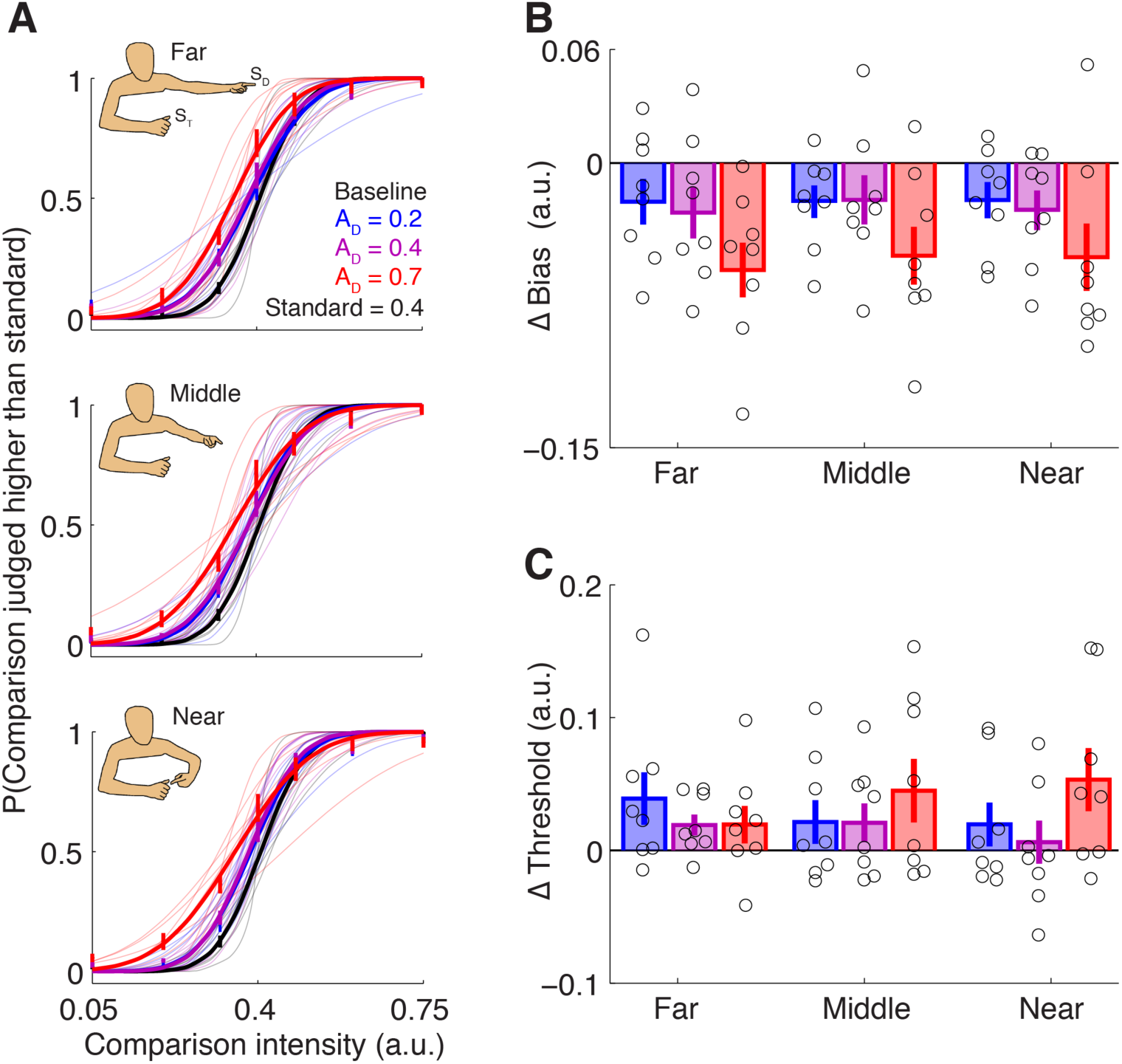
Bimanual interactions in intensity perception (conventions as in **Fig. 1**). (*n* = 8) (***A***) Average choice probability and psychometric curves from the group and individual subjects. Distractors attenuated the perceived intensity of the target stimulus depending on distractor amplitude (A_D_). Effects were similar across locations. (***B***) Changes in bias (PSE), with respective to baseline, with each distractor amplitude and location. Markers indicate individual subjects. Error bars indicate s.e.m. (***C***) Changes in discrimination threshold (JND) with respect to baseline.

Because tactile contextual modulation patterns were feature-specific, we evaluated bimanual cue combination models in the frequency and intensity domains separately; however, in both domains, we tested the hypothesis that contextual modulation involves divisive normalization. To capture contextual modulation in the frequency domain, which depended on both distractor frequency and distractor location, we implemented a model that comprised three key components (**Fig. 3*A***): A frequency-based modulation function that determined the variance of the target and distractor cue representations, a location-based weighting function that determined the cue weights, and a normalization term that rescaled the weights. The frequency-and location-based functions represent modulatory effects of feature-based attention and spatial attention, respectively. Accordingly, this model bears close resemblance to normalization models of attention in the visual domain (11). Our model fully recapitulated the observed bias and threshold changes (**Fig. 3*B,C***). On average, the model accounted for 94±1.1% of the variance in split-half cross-validation tests performed on each subject’s data (**Fig. 3*D***). Moreover, the model explained 96±1% of the variance in across-subject cross-validation tests (**Fig. 3*D***), approaching the noise ceiling (98±1%) which provides an estimate of the maximum achievable model performance given measurement noise and inter-subject reliability in our sample. Importantly, we performed quantitative comparisons of a large number of models (**Table S1**) that each assumed a linear combination of the target and distractor cue estimates, but that differed in their treatment of the frequency and position contextual manipulations. The model comprising frequency-based reliability modulation, location-based cue weighting, and normalization substantially outperformed the alternative models according to a number of metrics (**Table S2**) and was the most probable model given our data based on Akaike weights (**Fig. S3**).

**Figure 3.**
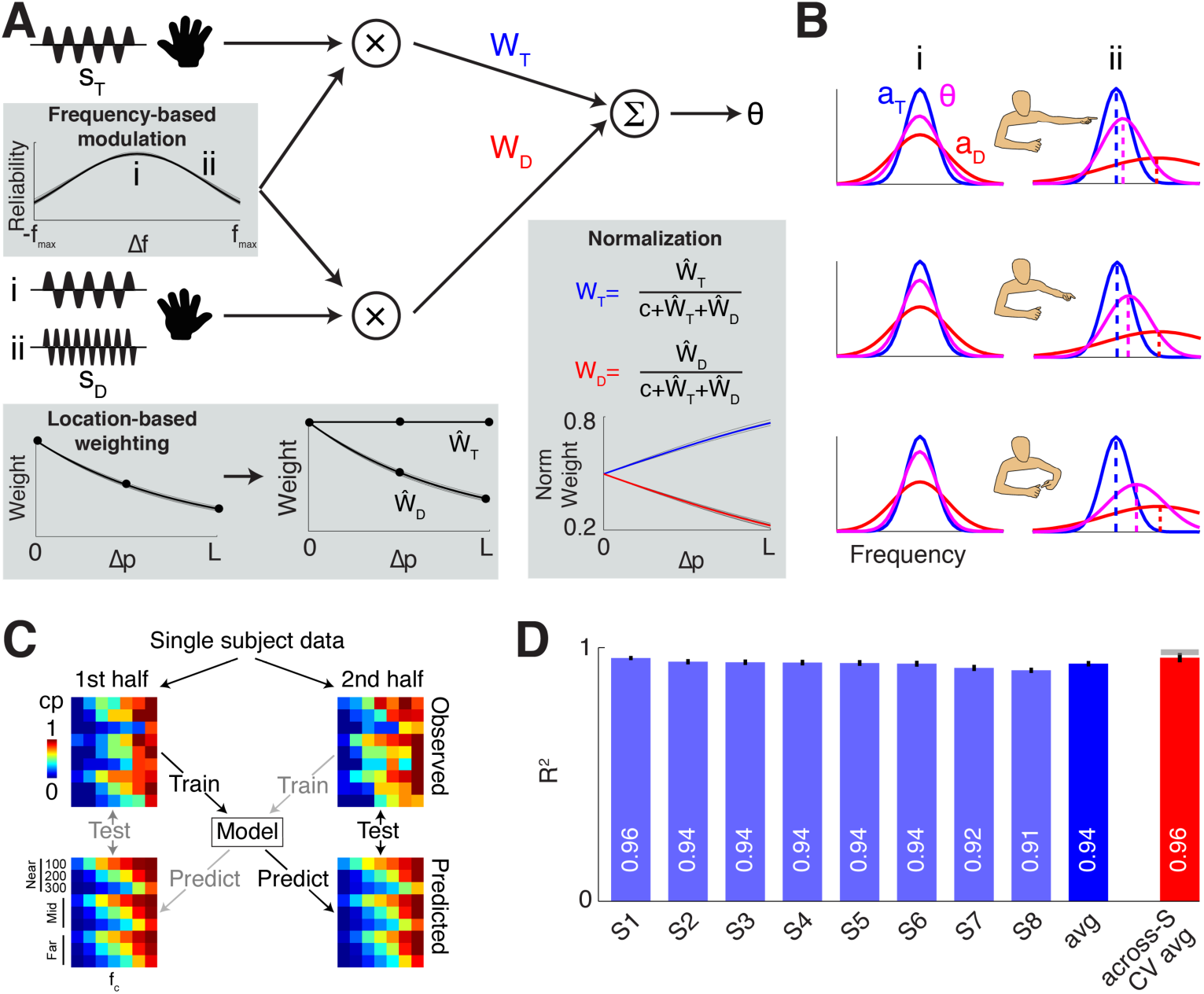
Model of tactile cue combination in the frequency domain. (***A***) An estimate of the target stimulus frequency (*θ*) is computed as a linear combination of the target (*s_T_*) and distractor stimulus cues (*s_D_*). Two example distractor frequencies, tested separately in the experiment, are denoted by *i* and *ii*. Cues are modeled by Gaussian probability distributions whose variances are initially set according to baseline discrimination thresholds. A model component capturing frequency-dependent effects (Frequency-based modulation) determines the modulated reliability of each cue according to the frequency disparity (*Δf*) between the cue and the attended (target) stimulus frequency. A model component capturing the position-dependent effects (Location-based weighting) determines each cue’s initial weight 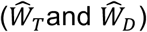 according to the position disparity (*Δp*) between the cue and the attended (target) stimulus position (*L* indicates the largest tested separation between the hands). Final target estimates are computed by linearly combining the attention-modulated cue representations using normalized weights *(W_T_* and *W_D_*). (***B***) Plots depict the probability distributions of the attention-modulated cues (*a_T_* and *a_D_*) and the final target estimate under a subset of example distractor conditions. Probability distributions are depicted for different distractor frequencies (columns; corresponding to *i* and *ii* from panel *a*) and relative hand positions (rows). (***C***) Split-half cross-validation was performed within each subject separately. Observed and predicted choice probability (CP) values are shown for an example subject. (***D***) Light and dark blue bars indicate single-subject and group-averaged cross-validated goodness-of-fit. Red bar indicates the average model performance in an across-subject cross-validation procedure. Horizontal gray bar indicates the noise ceiling. Error bars indicated s.e.m.

Contextual modulation in the intensity domain, which was invariant to hand location changes and was manifest in biasing effects only, could be explained with a simple normalization model (**Fig. 4*A***). The pooling of the target and distractor intensity representations for the normalizing factor explains why increases in distractor amplitude result in greater reductions in the perceived intensity of the target cue (**Fig. 4*B,C***). The normalization model accounted for 92±2% of the variance in within-subject cross-validation tests (**Fig. 4*D***). The normalization model also accounted for 94±2% of the variance in across-subject cross-validation tests (**Fig. 4*D***), which fell close to the theoretical maximum goodness-of-fit (95±3%). Because distractors only attenuated the perceived intensity of the target cue, their modulatory effects could not be explained by cue averaging (**Table S3**).

**Figure 4.**
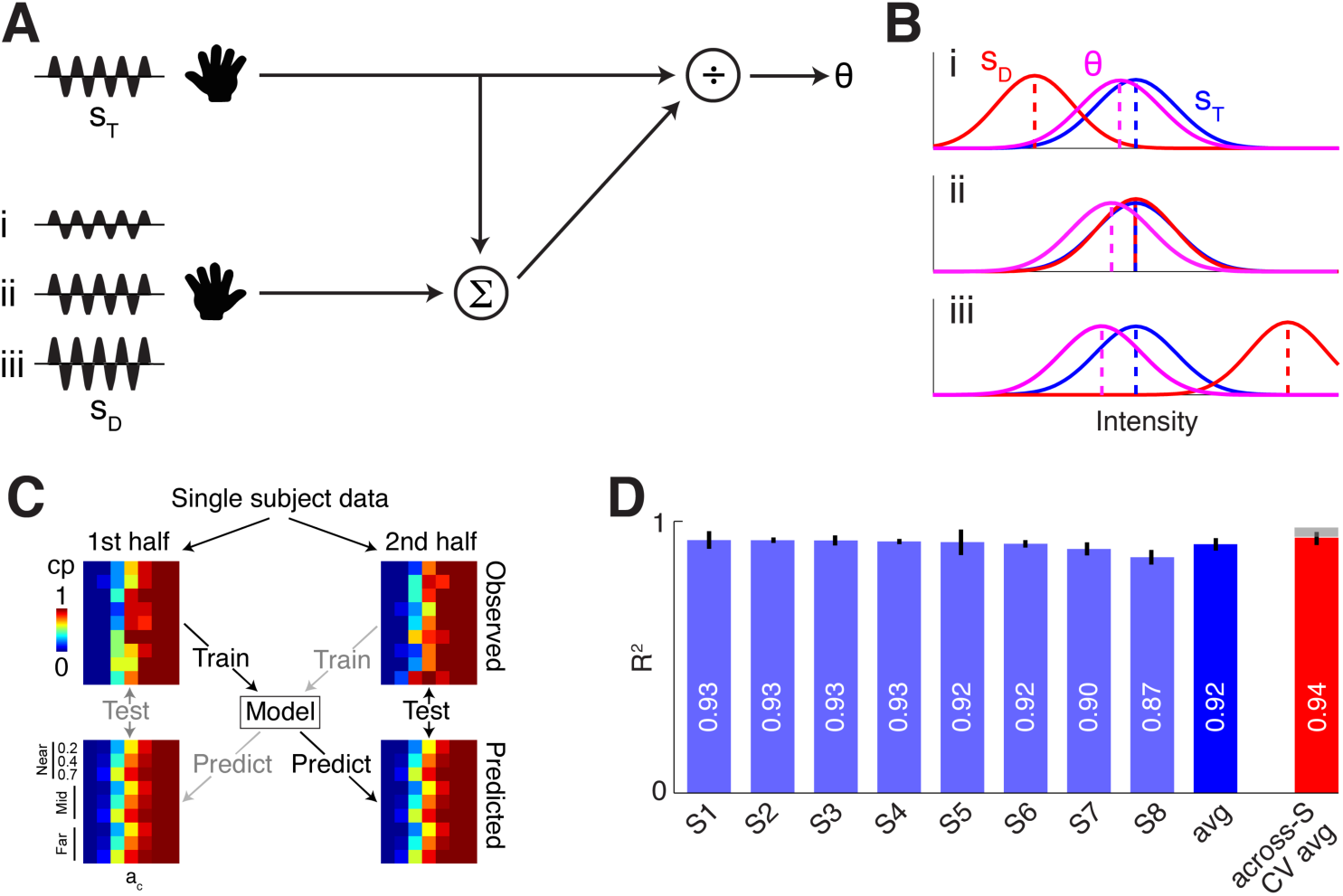
Model of tactile cue combination in the intensity domain. (***A***) An estimate of the target stimulus intensity (*θ*) is computed from the target (*s_T_*) and distractor stimulus cues (*s_D_*). Three distractor amplitudes, tested separately in the experiment, are denoted by *i, ii,* and *iii*. The target cue is normalized by the summed estimates of the cues to yield the final target intensity estimate. (***B***) Plots depict the probability distributions of the target and distractor cues (*s_T_* and *s_D_*) and the final target estimate under variations of distractor intensity (*i, ii*, and *iii* from panel *a*). (***C***) Split-half cross-validation was performed within each subject separately. Observed and predicted choice probability (CP) values are shown for an example subject. (***D***) Light and dark blue bars indicate single-subject and group-averaged cross-validated goodness-of-fit. Red bar indicates the average model performance in an across-subject cross-validation procedure. Horizontal gray bar indicates the noise ceiling. Error bars indicated s.e.m.

## Discussion

We found that contextual modulation in touch, probed through bimanual interactions, are characterized by idiosyncratic patterns that depend on whether attention is directed to the frequency or intensity of the vibration cues. Cue combination in the frequency domain is marked by attractive interactions that depend on the locations of the hands while interactions in the intensity domain only consist of attenuation which is invariant to hand location changes. Notably, our modeling results imply that divisive normalization, a canonical computation (5), is critical for tactile contextual modulation irrespective of the attended feature. These collective results provide clear demonstrations of how theoretical frameworks relating attention modulation and gain control can elucidate the processes by which touch experienced on one finger influences perception on another finger (10,12–16) and how cutaneous sensing can be modulated by proprioception (17–21).

That different patterns characterize bimanual interactions in the frequency and intensity domains is perhaps unsurprising given that these features are represented by distinct neural codes in somatosensory cortex (22). Normalization of rate-encoded intensity signals conceivably underlies the perceptual attenuation of the target cue intensity, as has been proposed for masking effects on tactile detection (10). The neural mechanisms underlying the representation and integration of vibration frequency remain more enigmatic. Our results suggest that the reliabilities of target and distractor frequency representations are modulated and differentially weighted before they are normalized and integrated. This contextual processing, which we interpret as reflecting a combination of attention modulation and normalization, may parallel the interactions between attention and divisive normalization previously described for visual processing (11). Because vibration frequency is represented by spike timing in cortical neuron populations that phase-lock to vibrations experienced on the skin (22–24), feature-based attention may enhance this temporal code’s precision (25,26). Alternatively, attention could modulate activity in putative downstream neurons that presumably convert the timing-based signals into rate-based representations (27,28). These representations could then be integrated through mechanisms analogous to those described for multisensory cue combination (29), which also involve normalization (30,31). To set the weights to the target and distractor cues, spatial attention could either directly modulate the activity representing the cues or attention could change the weighting of the connections between the populations signaling this information (32). Whether contextual modulation occurs at a particular level in the neuraxis is unclear: Bimanual interactions would appear to implicate the higher-order somatosensory regions that contain neurons with bilateral receptive fields (33), although activity in earlier processing levels could be modulated and normalized via feedback or lateral connections. Identifying the neural populations involved in tactile contextual modulation and testing these potential mechanisms will require neurophysiology recording experiments. These experiments may provide insights into why frequency processing is sensitive to spatial context while intensity processing is not. Future experiments can also establish whether the same contextual modulation principles apply to the processing of vibrations and tactile flutter signals, which may be represented by distinct neural codes (23,34).

Bimanual interactions that are sensitive to hand location likely involve conjunctive neural coding of cutaneous and proprioceptive signals (35). What reference frames or coordinate systems this conjunctively coded information occupies is an important consideration. Cutaneous events may be maintained in limb-based coordinates irrespective of their location in external space (36–38). In this case, tactile contextual modulation may be more probable and stronger when the hands are held in postures that dominate the statistics of bimanual actions or are learned through sensorimotor training (39,40). Alternatively, cue combination may occur after the tactile events are remapped from anatomical coordinates to external space (21,41–43). In this case, tactile contextual modulation may occur regardless of specific limb postures or even the contacted skin regions. Results from two control experiments support the remapping account. First, we observed robust proximity-dependent modulation patterns that were comparable in magnitude with the arms crossed and uncrossed (**Fig. S4**). Second, we found that distractors presented on the left forearm rather than the left hand also modulated frequency discrimination performance in a location-dependent manner (**Fig. S5**). These results implicate posterior parietal cortex in mediating tactile contextual interactions given its critical role in attention and coordinate transformations (41,44,45).

Our results reveal obligatory interactions between bimanual sensory cues, the patterns of which differ according to the attended stimulus feature. Despite these differences, which likely reflect distinct neural operations, tactile contextual modulation in both the frequency and intensity domains involve divisive normalization. These results reinforce the notion that normalization is a canonical neural computation which appears omnipresent in support of diverse functions like sensory processing, cue integration, attention, and decision-making (5). Our study may also have important clinical implications in light of recent efforts to relate divisive normalization to the atypical perceptual patterns associated with autism spectrum disorders (ASD) (46,47). Indeed, the prevalence of aberrant somatosensory processing is high in individuals with ASD and other neurodevelopmental disorders (48–50), and our study offers a tractable paradigm for exploring potential links between aberrant touch and alterations in contextual modulation. Future studies with neurotypical and clinical populations must also address how bimanual contextual modulation interacts with and supports bimanual motor control (51,52).

## Materials and Methods

### Participants

A total of 30 subjects participated in at least 1 of 5 experiments. Eight subjects (4m4f; mean age ± standard deviation: 24.5 ± 8.6 years; mean arm length ± standard deviation: 70.8 ± 4.9 cm) performed the main frequency discrimination task (Experiment 1). Eight subjects (3m5f; 22 ± 5.3 years; 67 ± 3.6 cm) performed the main intensity discrimination task (Experiment 2). Eight subjects (3m5f; 21.56 ± 3.9 years; 65.8 ± 3.2 cm) performed the frequency discrimination task in a control experiment with the arm crossing manipulation (Experiment 3). Eight subjects (3m5f; 25.4 ± 5.3 years; 66 ± 3.5 cm) performed the frequency discrimination task in a control experiment in which distractors were delivered to the left forearm (Experiment 4). Eight subjects (2m6f; 22.8 ± 4.6 years; 65.5 ± 3.6 cm) performed the frequency discrimination task on the right hand in a control experiment in which the position of the left hand was manipulated without distractor stimulation (Experiment 5). One subject participated in Experiments 3, 4, and 5; one participated in Experiments 1 and 5; one participated in Experiments 2 and 3; one participated in Experiments 3 and 5; two subjects participated in Experiments 1 and 4; and two subjects participated in Experiments 4 and 5. All subjects were right-handed according to the Edinburgh Handedness Inventory (53) and reported normal somatosensory functions. Testing procedures were performed in compliance with the policies and procedures of the Baylor College of Medicine Institutional Review Board. All participants provided informed written consent and were paid for their participation or declined compensation.

### Experiment 1. Frequency discrimination task

Tactile frequency discrimination was tested using a two-alternative forced choice (2AFC) design. On each trial, participants experienced vibrotactile stimulation on their right thumb (target digit, D_T_) in two intervals: One interval (randomized over trials) always contained the 200-Hz standard stimulus and the other contained a comparison stimulus whose frequency varied (100, 140, 180, 200, 220, 260, 300Hz). Stimuli were delivered at amplitudes that were equated for perceived intensity (54). Subjects reported which interval contained the stimulus perceived to be higher in frequency. Participants performed the task in the absence (baseline) and presence of vibrotactile distractors presented to their left index finger (distractor digit, D_D_). Subjects were explicitly instructed to ignore the distractor stimuli. On each trial, distractors paired with the comparison stimulus were always matched in frequency. Distractors paired with the 200-Hz standard stimulus were set at 100, 200, or 300Hz. This design enabled us to characterize frequency-dependent distractor influences on the perception of the 200-Hz standard stimulus.

We also manipulated the position of D_D_ to test whether distractor influences depended on the proximity of the hands. D_T_ was always positioned 15cm in front the participant’s body (midline) with the finger pad facing away from the body. D_D_ was positioned at three locations (**Fig. S6A**): D_D_ was 1cm from D_T_ (near position), D_D_ was co-planar and aligned with D_T_ as the left arm was fully extended and angled 60° with respect to the body (far position), or D_D_ was located at the midpoint between the near and far positions (middle position). Magnetic holders secured the fingers to the table surface ensuring that hand positions and alignments were consistent throughout the experiment. Participants’ hands were visually occluded using custom-designed goggles.

Each subject was tested in 2 sessions (inter-session interval: 4.3 ± 4.6 days). Within a session, an equal number of trials were tested under the baseline condition (D_T_ with no distractor), 9 distractor conditions (3 distractor frequencies x 3 distractor positions), and a condition in which the discrimination task was performed using D_D_ only. In each condition, the total repetitions of each comparison frequency depended on its absolute difference with respect to 200Hz (Δ100Hz = 10 reps; Δ60Hz = 11 reps; ≤Δ20Hz = 12 reps) to bias sampling to the more challenging comparisons. This yielded 78 trials per condition and 858 trials per session. In each session, baseline trials were randomized and tested in a single block. Trials involving D_D_ only were similarly tested. Distractor trials were pseudo-randomized according to distractor position. Each distractor position was tested in 3 blocks yielding a total of 9 distractor blocks, the order of which was counterbalanced over subjects and sessions. Subjects were provided 2-3 min between blocks to rest. Over two sessions, this study design yielded 1716 trials per subject.

### Experiment 2. Intensity discrimination task

Subjects’ ability to discriminate vibration intensity was tested using the same 2AFC procedure as described for Experiment 1. On each trial, participants judged the intensity of tactile stimulus pairs that were delivered to their right thumb. All stimuli were matched in frequency (200Hz) and were clearly suprathreshold. The amplitude of the standard stimulus was 0.4 (arbitrary units). The amplitude of the comparison stimuli varied (0.05, 0.2, 0.32, 0.4, 0.48, 0.6, 0.75). The voltage measured from the amplifier’s output ranged from 0.6–3.3V over the tested vibration amplitudes. Subjects performed the intensity discrimination task in the absence and presence of distractors. The amplitude of the distractors, presented to the left index finger, were either 0.2, 0.4, or 0.7. As in Experiment 1, distractor stimuli presented during the comparison stimulus interval always matched the target stimulus experienced on D_T_. The position of the distractor digit was manipulated as in Experiment 1. Over two sessions, this study design yielded 1716 trials per subject.

### Experiment 3. Frequency discrimination task with arm crossing manipulation

The procedure was similar to that described for Experiment 1 with the major exception that only two limb configurations were tested: Arms uncrossed and crossed (**Fig. S4** and **Fig. S6B**). In the uncrossed configuration, the target digit (right thumb) and distractor digit (left index finger) were positioned 15cm in front of the body to the right and left of midline, respectively. In the crossed configuration, the locations of the hands were reversed such that the right and left hands were positioned to the left and right of midline, respectively. In both configurations, the distance between the target and distractor digits was 20cm. Two distractor frequencies (100 and 300Hz) were tested. This design enabled us to test whether bimanual interactions in the frequency domain were associated with specific limb configurations or simply the distance between the hands irrespective of limb configuration. Over two sessions, this study design yielded 936 trials per subject.

### Experiment 4. Frequency discrimination task with forearm distractors

The procedure was similar to that described in Experiment 1 with the major exception that the distractors were presented on the left forearm rather than the left hand. As in Experiment 1, the target digit was maintained in front of the body at midline and the left arm was repositioned to manipulate the location of the distractor (**Fig. S5** and **Fig. S6C**). In the near position, the stimulated forearm site was 1cm from the target digit. In the far position, the left arm was extended away from the target digit. Two distractor frequencies (100 and 300Hz) were tested. This design enabled us to test whether position-dependent cutaneous interactions in the frequency domain only occur under bimanual stimulation conditions. Over two sessions, this study design yielded 780 trials per subject.

### Experiment 5. Frequency discrimination task with manipulations of the distractor hand in the absence of distractor stimulation

The procedure was identical to that described for Experiment 1 except that no distractor stimulation was presented. In this experiment, we only manipulated the position of the left hand (**Fig. S2** and **Fig. S6a**) while subjects performed the frequency discrimination task using their right hand. In a single session, this study design yielded 432 trials per subject.

### Vibrotactile stimulation

Sinusoidal signals were digitally generated (sampling rate: 44.1kHz) in Matlab (2011b, MathWorks) running on a Macbook Pro (model A1278; OS X 10.9.5, 2.5GHz Core i5, 4GB RAM)(54). The experiment was controlled using Psychtoolbox-3. Each 800-ms stimulus was characterized by linear on- and off-ramps (ramp duration: 50ms). Stimuli on each trial were separated by an 800-ms inter-stimulus interval. The signals were outputted via the auxiliary port, amplified (Krohn-Hite Wideband Power Amplifier, model 7500), and delivered to the skin through miniature electromechanical tactors (Fingers: type C-F, Engineering Acoustics, Inc.; Forearm: type C-2, Engineering Acoustics, Inc.). Tactors were fastened to the distal phalanges of the index fingers or the left forearm self-adherent cohesive wrap bandages. Subjects wore earmuffs (Peltor H10A Optime 105 Earmuff, 3M) to attenuate any sounds associated with the tactile stimulation.

### Data Analysis

Analyses were performed using Matlab and R-studio. To quantify each participant’s ability to discriminate tactile frequency in the baseline and distractor conditions, we fitted each subject’s performance data with a Gaussian cumulative distribution function:

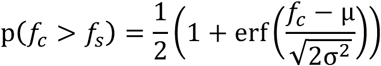

where *p(f_c_ > f_s_*) is the choice probability (CP) indicating the proportion of trials a comparison stimulus with frequency *f_c_* was judged to be higher in frequency than the standard stimulus *f_s_, μ* and *σ* are free parameters corresponding to the point of subjective equality (PSE) and just-noticeable difference (JND), respectively, and erf(*x*) is the error function of *x*. The PSE is a measure of bias and indicates the comparison frequency perceived as equal to the standard frequency. The JND is a measure of sensitivity that is defined as the standard deviation of the Gaussian, which corresponds to 84% performance. Participants’ ability to discriminate vibration intensity in Experiments 2 was also quantified using the Gaussian cdf.

In group-level analysis, we determined whether baseline-subtracted PSE and JND estimates differed significantly according to distractor conditions. In Experiments 1–4, we conducted twoway repeated-measures ANOVA (rmANOVA) with frequency (or intensity) and position as the within-subjects factors. In Experiment 5, we conducted a one-way rmANOVA with distractor hand position as the within-subjects factor.

### Modeling tactile cue combination

#### General

To understand the computing principles underlying tactile cue combination, we implemented and compared competing models for bimanual interactions in the frequency and intensity domains separately. For each model, we assumed that the nervous system initially represents the target stimulus and distractor stimulus as Gaussian probability distributions 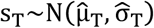 and 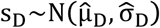, respectively. Based on these representations, we assumed that the nervous system computes a final estimate of the target stimulus, θ~N(μ_θ_, σ_θ_). The variances of the initial target and distractor representations were determined empirically in the psychophysical experiments.

#### Cue combination in the frequency domain

We assumed that the final target estimate is computed as a weighted combination of the target and distractor representations. We also assumed that the weighting of the target and distractor representations as well as their variances (reliability) can be modulated before the final target estimate is computed. Thus, the model takes the general form:

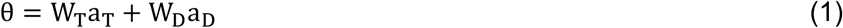

where, θ is the representation of the final estimate, a_T_~N (μ_T_, σ_T_) and a_D_~N (μ_D_, σ_D_) are the modulated representations of s_T_ and s_D_, respectively, and W_T_ and W_D_ are their respective weights. According to this model, the predicted bias (μ_θ_, PSE) and threshold (σ_θ_, JND) of the final estimate are determined as:

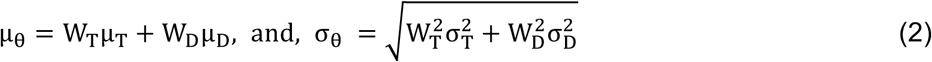

The most probable model given our data (see *Model selection*) comprised 4 free parameters and included three key functions: frequency-based cue reliability modulation, location-based cue weighting, and normalization. The frequency-based reliability modulation function sets the reliability of each cue representation according to the cue’s frequency:

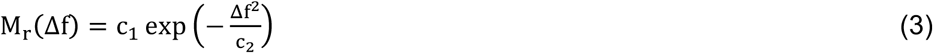

where the modulation values in M_r_ are normally distributed over Δf (the difference in frequency between each cue and the standard frequency), and c_1_ and c_2_ are free parameters. This component can be considered to reflect feature-based attention directed toward the standard frequency, 200Hz. The function determines the modulated reliabilities of both the target and distractor cues:

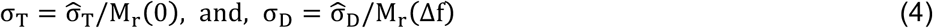

Although the modulation function’s impact is substantially greater on the distractor representations, the function causes a slight increase in the variance of the target cue representation (Fig. 3b) which is consistent with an attentional cost to ignoring the distractor. Also, because the modulation function only sets the variance of the cue representations, 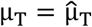 and 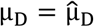.

The location-based weighting function determines the contribution of each cue according to the cue’s position:

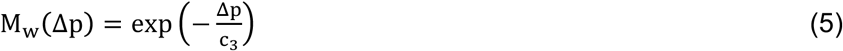

where M_w_ is exponentially distributed over Δp (the distance between the target and distractor hands) and c_3_ is a free parameter. To account for variations in subject arm lengths, the relative distances in the near, middle, and far positions were 0, 0.5, and 1, respectively. This component can be considered to reflect spatial attention directed toward the location of the target hand. Accordingly, before normalization, the target cue is assigned the maximum weight by M_w_ under all of the limb configurations while the weight to the distractor cue decreases with greater inter-manual separation (Fig. 3a).

The final weights to the target (W_T_) and the distractor (W_D_) that determine their contributions to the final estimate are computed using a normalization function that includes a single free parameter, c_4_:

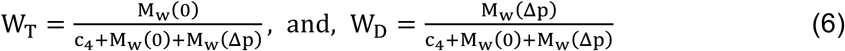

We tested a total of 19 alternative models. All of the models assumed a linear combination of the target and distractor representations (Eqs. 1, 6) but differed in the determination of the target and distractor cue reliabilities and their weights (**Table S1**).

#### Cue combination in the intensity domain

We separately modeled distractor influences on tactile intensity perception, which were limited to changes in bias. The most probable model given our data was one incorporating only divisive normalization: The perceived intensity of the target stimulus (μ_θ_) is estimated by normalizing the representation of the target cue amplitude with the sum of the target and distractor cue amplitudes:

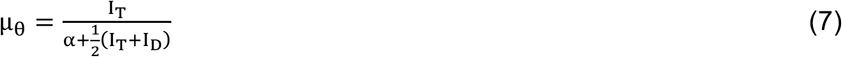

where I_T_ is the target amplitude, I_D_ is the distractor amplitude, and α is a free parameter. This model assumes no effects on discrimination thresholds in the intensity domain, so the threshold associated with the final estimate matches the threshold associated with the target cue alone, σ_θ_ = σ_T_.

We compared the normalization model to an alternative model based on simple cue averaging. In this model, the perceived intensity of the target stimulus is estimated by averaging the amplitudes of the target and distractor cues, again assuming no changes in the perceptual threshold:

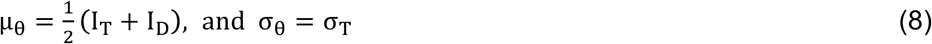

### Model fitting and performance assessment

#### Single subject fitting

Model parameters were estimated using a within-subject cross-validation procedure. Each subject’s data were divided into two sets that contained an equal number of repetitions for each experimental condition. This pseudo-random data partitioning permitted the calculation of unbiased choice probability values for each split-half dataset. Using two-fold cross-validation, the model was trained on one dataset using maximum likelihood estimation and tested on the second dataset. The squared correlation between the observed and predicted CP was computed as a measure of variance explained for each fold and the variance explained was averaged over both folds for a single goodness-of-fit measure. This two-fold cross-validation procedure was repeated 100 times for each subject.

#### Group-level fitting

To compare parameter estimates and model performance at a group level, we adopted a leave-one-subject-out cross-validation approach. On each fold, the model was trained on the full datasets of 7 subjects and tested on the data of the held-out subject. The goodness-of-fit calculated in this manner provides an estimate of the inter-subject reliability of the models. Using the same leave-one-subject-out procedure, we calculated the response variance in a single subject explained simply by the averaged data over the other 7 subjects. The group-averaged data provided a parameter-free model which established a noise ceiling against which we compared the goodness-of-fit values calculated for the fitted models.

We additionally conducted Bayesian analyses to evaluate each candidate model against the parameter-free model established by the leave-one-subject-out cross-validation (**Table S2** and **Table S3**). For each model separately, we quantified the relative support for the alternative hypothesis that the model was outperformed by the parameter-free model (H_1_) compared to the null hypothesis of no performance differences (H_0_) where the Bayes factor (BF) was p(data|H_1_)/p(data|H_0_). Thus, smaller BF values indicated stronger evidence for each fitted model.

### Model selection criteria

We adopted a number of complementary approaches for selecting a favored model among the alternative models.

#### Log-likelihood

If an experiment consists of n independent Bernoulli trials, each having probability of success (p) and total number of successes in the trials (x) then the likelihood is:

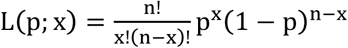

Maximizing the likelihood is equivalent to maximizing the log-likelihood (ll(p;x)), which is given by:

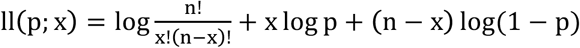

The first term of the above equation is a constant that does not involve the parameter p, where p is expressed as a Gaussian cdf. So, for a given model, we maximize:

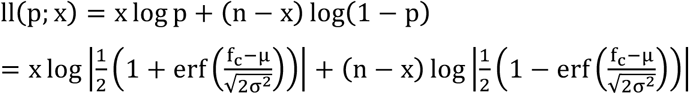

where, μ and σ are the bias and threshold parameters, respectively, estimated for the CP values predicted by the model. The log-likelihood value associated with each model was computed and we favored the model identified with the maximum log-likelihood value.

#### Residual Sum of Squares (RSS)

We computed RSS, the sum of squared errors between the measured data (p) and the data predicted 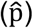 by the model, as a measure of prediction error. We favored the model with the minimum RSS value.

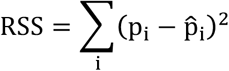

#### Akaike Information Criteria (AIC)

We computed AIC as a selection measure that balanced model complexity (k, number of estimated parameters) and model performance (L, the maximum likelihood of the model). We favored the model with the lowest AIC value.

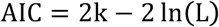

#### Bayesian Information Criteria (BIC)

We computed BIC as another selection measure that balanced model complexity and performance while adjusting for the number of fitted data points (n). As with AIC, we favored the model with the lowest BIC value.

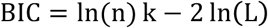

#### Likelihood Ratio Test (LRT)

We performed LRT calculations of pairwise model comparisons: One model was designated the null model and the other was designated the alternative model.

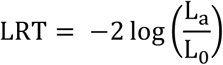

where L_0_ is the likelihood of the observations under the null model and L_a_ is the likelihood of the observations under the alternative model. We favored the alternative model if LRT was positive and the null model if LRT was negative.

#### Probability or Akaike weights

Akaike weights are estimated by normalizing the model likelihood such that the relative likelihoods for all models under consideration sum to 1. Hence, these weights can be considered as reflecting the probability or “weight of evidence” in favor of a given model in a fixed set of alternative models (55). The weight w_i_ of the i^th^ model is computed as:

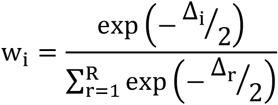

where Δ_i_ = AIC_i_ – AIC_min_ and AIC_min_ is the minimum of the AIC_i_ values over the R alternative models. We favored the model with the highest Akaike weight.

## Data and Code Availability

The behavioral data (.mat files) and analysis code are available at https://github.com/YauLab/PROPCUT_MSR2018.

## Author contributions

MSR and JMY designed the experiments, MSR collected and analyzed the data, MSR and JMY wrote the manuscript.

## Acknowledgements

This work was performed in the Neuromodulation and Behavioral Testing Facilities of BCM’s Core for Advanced MRI (CAMRI). We thank Akshat Patel, Sriparna Sen, and Silvia Convento for help with data collection and Umme Farjana for help with graphics editing. We thank Yau Lab members for helpful discussions and comments on early manuscript drafts. This work was supported by BCM seed funds and the Alfred P. Sloan Research Fellowship.

## Supplemental Information

**Supplemental Figure 1.**
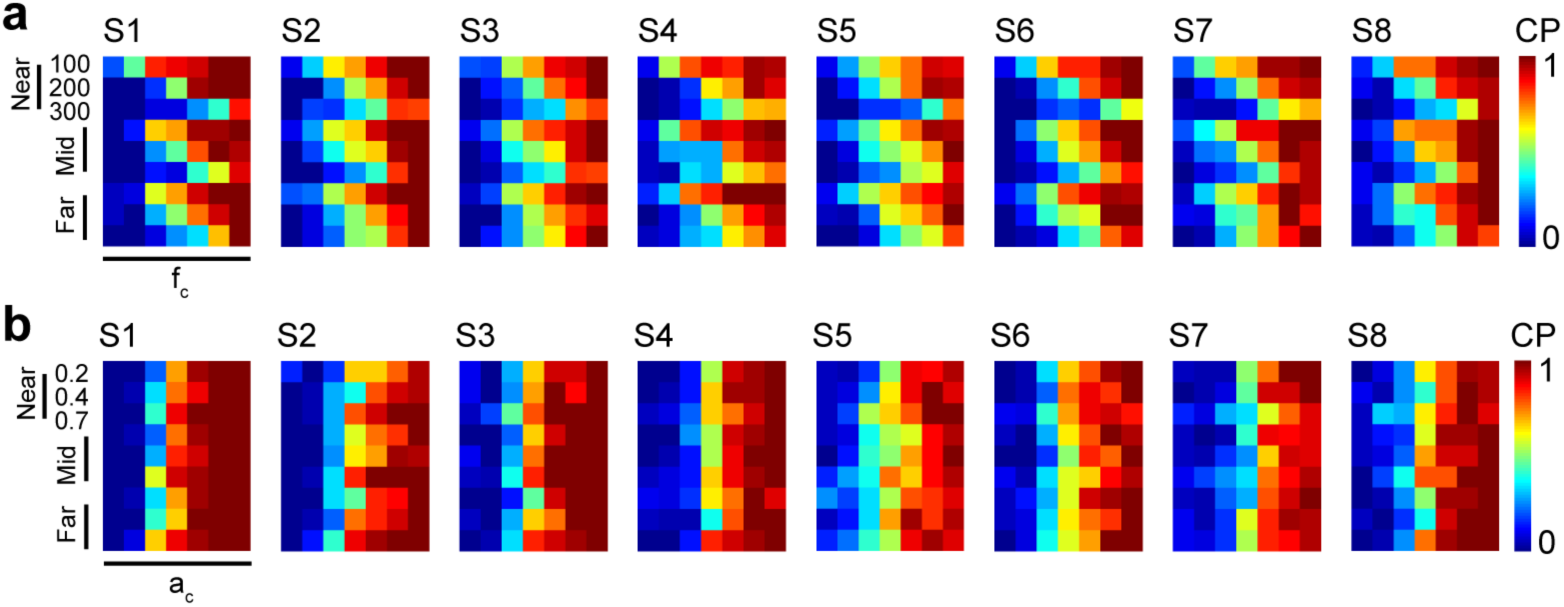
Choice probability (CP) for individual subjects in Experiments 1 and 2. Each plot indicates data from a single subject. (**a**) Heat maps indicate CP for frequency discrimination task in Experiment 1. In each plot, columns correspond to comparison frequencies (f_c_) and rows correspond to distractor conditions (distractor frequencies and locations). (**b**) CP for intensity discrimination task in Experiment 2. In each plot, columns correspond to comparison amplitudes (a_c_) and rows correspond to distractor conditions (distractor amplitudes and locations). Subjects who participated in Experiment 2 did not necessarily participate in Experiment 1.

**Supplemental Figure 2.**
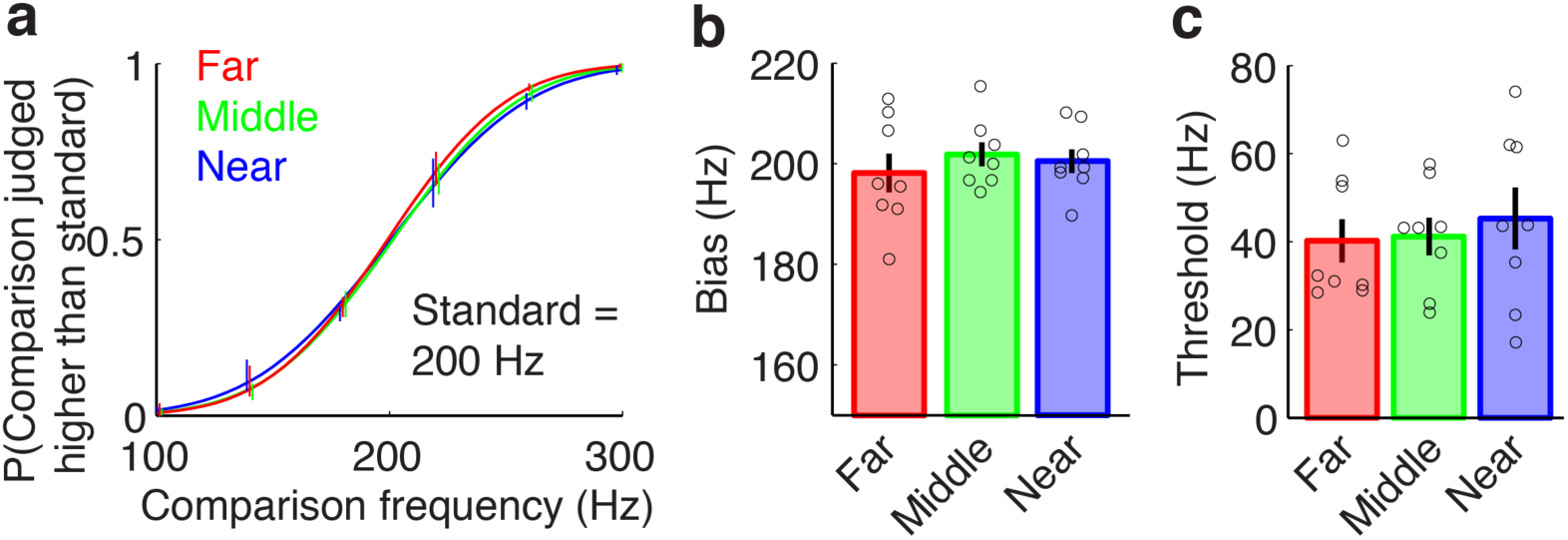
Effect of the distractor hand location on the frequency discrimination using the target hand (Experiment 5, n=8). (**a**) Group-averaged choice probability values and psychometric curves in the far, middle and near distractor locations. (**b**) Bias estimates did not differ significantly across hand locations (one-way rmANOVA, F(2,14)=0.33, p=0.73, η_p_^2^=0.04). (**c**) Threshold estimates did not differ significantly across hand locations (one-way rmANOVA, F(2,14)=0.81, p=0.47, η_p_^2^=0.1). Markers indicate individual subject parameter estimates. Error bars indicate s.e.m.

**Supplemental Figure 3.**
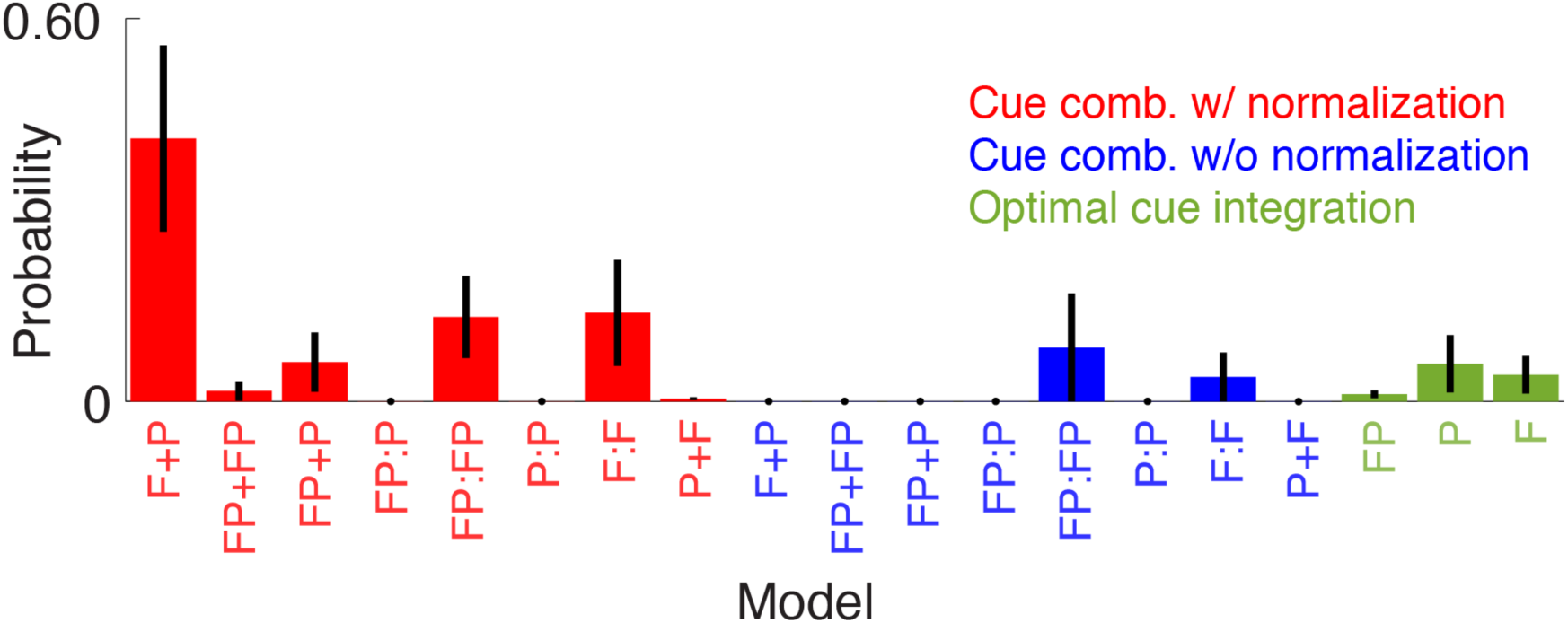
Relative probabilities of frequency combination models. Three general model classes were tested (also see **Supplemental Table 1**): Cue combination with weight normalization (red), cue combination without weight normalization (blue), and optimal cue integration (green). Models within a class varied by their cue reliability modulation and weighting functions. Each model’s relative probability was computed with respect to all other model Akaike information criterion values (**Supplemental Table 2**). Across models fit to each subject individually (Experiment 1, n=8), the model comprising frequency-based cue reliability modulation, location-based cue weighting, and normalization was the most probable of all. Error bars indicate s.e.m.

**Supplemental Figure 4.**
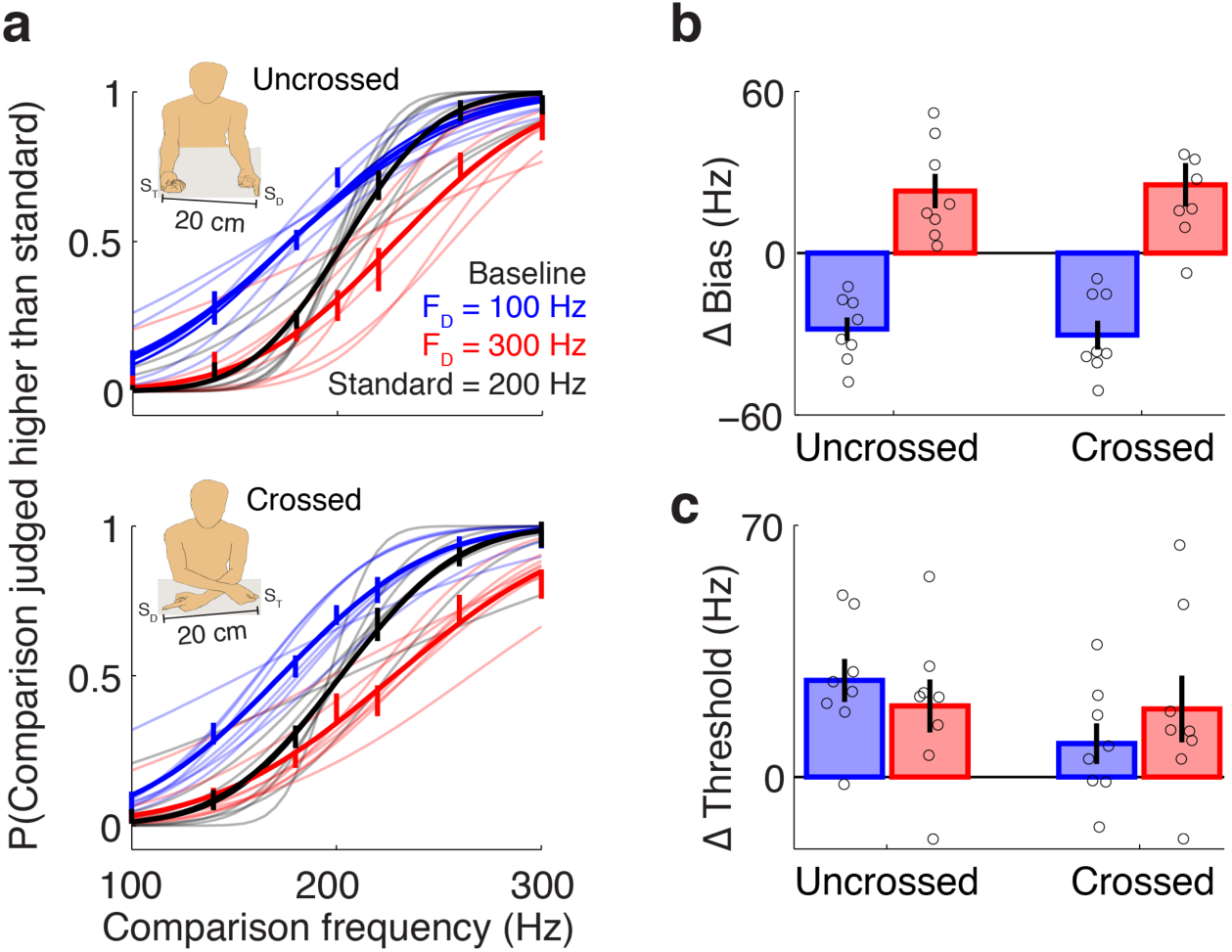
Influence of arm crossing on multi-finger interactions in the frequency domain (Experiment 3, n=8). (**a**) Distractors exert frequency-dependent changes to tactile discrimination performance in the uncrossed and crossed limb postures. The absolute separation between the hands was matched in the two postures. Group-averaged (thick traces) and individual subject psychometric curves (thin traces) are shown. (**b**) Frequency-specific distractor effects on PSE did not differ between the uncrossed and crossed postures (two-way rmANOVA, distractor frequency main effect: F(1,7)=265.7, p=8e-7, η_p_^2^=0.97; limb posture main effect: F(1,7)=0.1, p=0.77, η_p_^2^=0.01; interaction effect: F(1,7)=0.23, p=0.65, η_p_^2^=0.03). (**c**) Distractor effects on JND did not differ significantly according to distractor frequency or limb posture (two-way rmANOVA, distractor frequency main effect: F(1,7)=0.06, p=0.81, η_p_^2^=0.01; limb posture main effect: F(1,7)=1.1, p=0.34, η_p_^2^=0.13; interaction effect: F(1,7)=3.5, p=0.10, η_p_^2^=0.33). Markers indicate individual subject parameter estimates. Error bars indicate s.e.m.

**Supplemental Figure 5.**
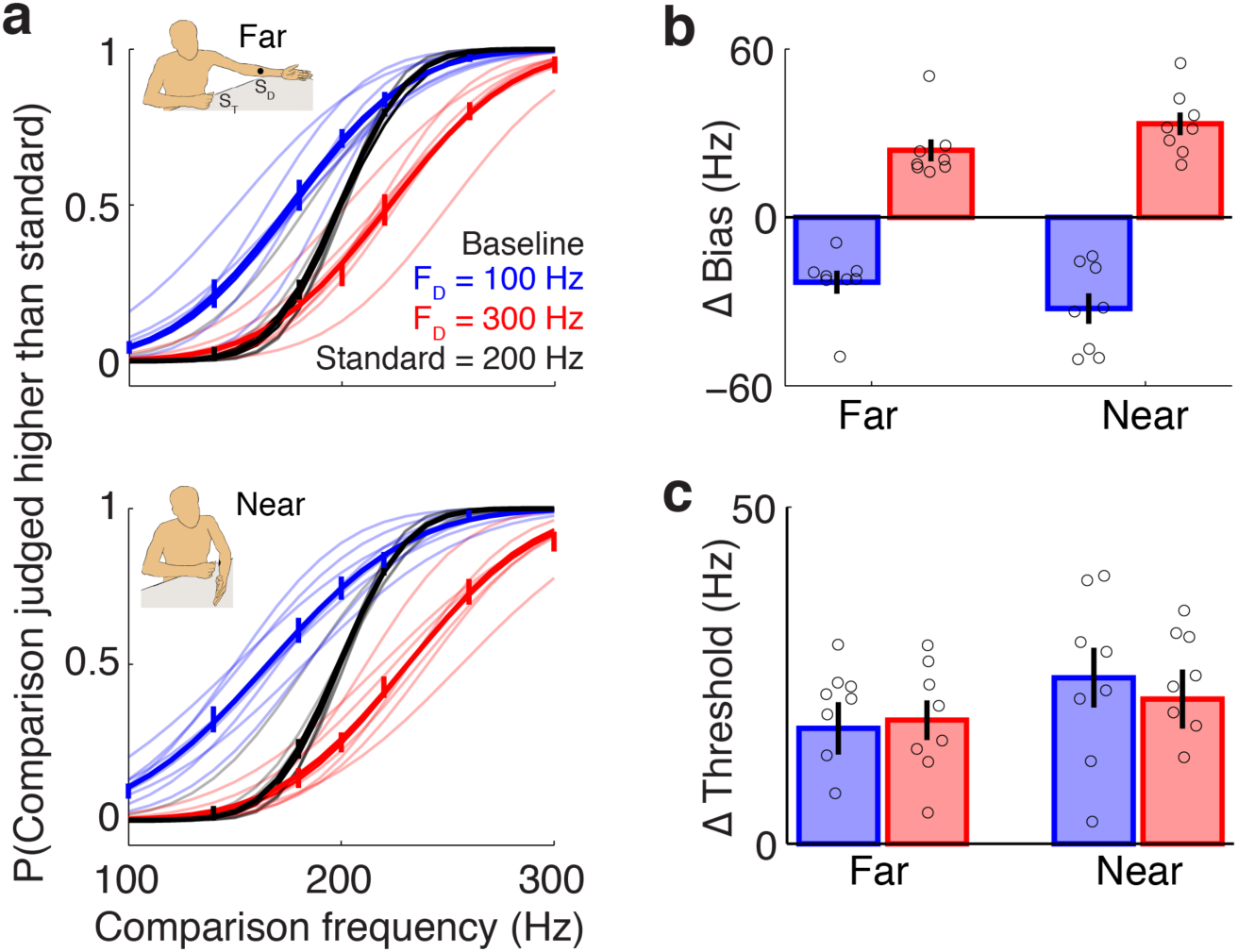
Distractors experienced on the forearm influence frequency discrimination performance in a frequency- and position-dependent manner (Experiment 4, n=8). (**a**) Distractors exert frequency-dependent influences on tactile discrimination performance in the far and near distractor positions. Group-averaged (thick traces) and individual subject psychometric curves (thin traces) are shown. **b**) Frequency-specific distractor effects on PSE depended on the proximity of the contacted sites (two-way rmANOVA, distractor frequency main effect: F(1,7)=154.8, p=5e-6, η_p_^2^=0.96; distance main effect: F(1,7)=0.0001, p=0.99, η_p_^2^=0; interaction effect: F(1,7)=10.1, p=0.02, η_p_^2^=0.59). (**c**) Distractor effects on JND depended on the proximity of the contacted sites (two-way rmANOVA, distractor frequency main effect: F(1,7)=0.09, p=0.77, η_p_^2^=0.01; distance main effect: F(1,7)=7.2, p=0.03, η_p_^2^=0.51; interaction effect: F(1,7)=0.43, p=0.53, η_p_^2^=0.06). Markers indicate individual subject parameter estimates. Error bars indicate s.e.m.

**Supplemental Figure 6.**
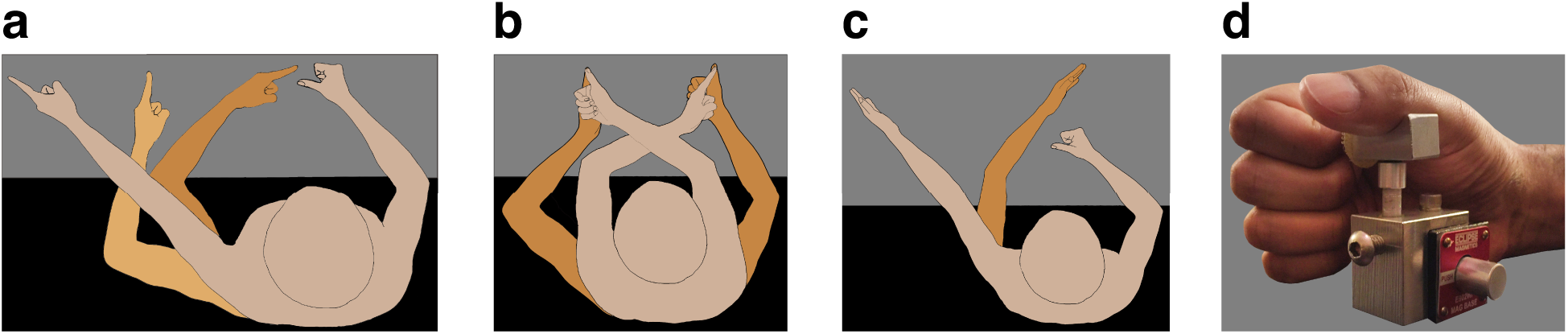
Experimental setup. (**a**) Hand positions (far, middle, and near conditions) tested in Experiments 1, 2, and 5. (**b**) Crossed and uncrossed conditions tested in Experiment 3. (**c**) Limb postures tested for forearm-finger interactions in Experiment 4 (near and far conditions). (**d**) Stimulated fingers were supported by finger-holders to ensure reliable positioning throughout the experiment.

**Supplemental Table 1.** Frequency combination models

| Model | | $M_r$ | $\sigma_T$ | $\sigma_D$ | $M_w$ | $W_T$ | $W_D$ |
| --- | --- | --- | --- | --- | --- | --- | --- |
| Normalization | F+P | $M_r(\Delta f)$<br>$= c_1 \exp\left(-\frac{\Delta f^2}{c_2}\right)$ | $\frac{\hat{\sigma}_T}{M_r(0)}$ | $\frac{\hat{\sigma}_D}{M_r(\Delta f)}$ | $M_w(\Delta p)$<br>$= \exp\left(-\frac{\Delta p}{c_3}\right)$ | $\frac{M_w(0)}{c_4 + M_w(0) + M_w(\Delta p)}$ | $\frac{M_w(\Delta p)}{c_4 + M_w(0) + M_w(\Delta p)}$ |
| | FP+FP+N | $M_r(\Delta f, \Delta p) =$<br>$c_1 \exp\left[-\left(\frac{\Delta f^2}{c_2} + \frac{\Delta p}{c_3}\right)\right]$ | $\frac{\hat{\sigma}_T}{M_r(0,0)}$ | $\frac{\hat{\sigma}_D}{M_r(\Delta f, \Delta p)}$ | $M_w(\Delta f, \Delta p) =$<br>$\exp\left[-\left(\frac{\Delta f^2}{c_4} + \frac{\Delta p}{c_5}\right)\right]$ | $\frac{M_w(0,0)}{c_7 + M_w(0,0) + M_w(\Delta f, \Delta p)}$ | $\frac{M_w(\Delta f, \Delta p)}{c_7 + M_w(0,0) + M_w(\Delta f, \Delta p)}$ |
| | FP+P+N | $M_r(\Delta f, \Delta p) =$<br>$c_1 \exp\left[-\left(\frac{\Delta f^2}{c_2} + \frac{\Delta p}{c_3}\right)\right]$ | $\frac{\hat{\sigma}_T}{M_r(0,0)}$ | $\frac{\hat{\sigma}_D}{M_r(\Delta f, \Delta p)}$ | $M_w(\Delta p)$<br>$= \exp\left(-\frac{\Delta p}{c_4}\right)$ | $\frac{M_w(0)}{c_6 + M_w(0) + M_w(\Delta p)}$ | $\frac{M_w(\Delta p)}{c_6 + M_w(0) + M_w(\Delta p)}$ |
| | FP:P+N | $M_r(\Delta f, \Delta p) =$<br>$c_1 \exp\left[-\left(\frac{\Delta f^2}{c_2} + \frac{\Delta p}{c_3}\right)\right]$ | $\frac{\hat{\sigma}_T}{M_r(0,0)}$ | $\frac{\hat{\sigma}_D}{M_r(\Delta f, \Delta p)}$ | $M_w(\Delta p)$<br>$= \exp\left(-\frac{\Delta p}{c_3}\right)$ | $\frac{M_w(0)}{c_5 + M_w(0) + M_w(\Delta p)}$ | $\frac{M_w(\Delta p)}{c_5 + M_w(0) + M_w(\Delta p)}$ |
| | FP:FP+N | $M_r(\Delta f, \Delta p) =$<br>$c_1 \exp\left[-\left(\frac{\Delta f^2}{c_2} + \frac{\Delta p}{c_3}\right)\right]$ | $\frac{\hat{\sigma}_T}{M_r(0,0)}$ | $\frac{\hat{\sigma}_D}{M_r(\Delta f, \Delta p)}$ | $M_w(\Delta f, \Delta p)$<br>$= M_r(\Delta f, \Delta p)$ | $\frac{M_w(0,0)}{c_4 + M_w(0,0) + M_w(\Delta f, \Delta p)}$ | $\frac{M_w(\Delta f, \Delta p)}{c_4 + M_w(0,0) + M_w(\Delta f, \Delta p)}$ |
| | P+N | $M_w(\Delta p)$<br>$= c_1 \exp\left(-\frac{\Delta p}{c_2}\right)$ | $\frac{\hat{\sigma}_T}{M_w(0)}$ | $\frac{\hat{\sigma}_D}{M_w(\Delta p)}$ | NA | $\frac{M_w(0)}{c_3 + M_w(0) + M_w(\Delta p)}$ | $\frac{M_w(\Delta p)}{c_3 + M_w(0) + M_w(\Delta p)}$ |
| | F+N | $M_r(\Delta f)$<br>$= c_1 \exp\left(-\frac{\Delta f^2}{c_2}\right)$ | $\frac{\hat{\sigma}_T}{M_r(0)}$ | $\frac{\hat{\sigma}_D}{M_r(\Delta f)}$ | NA | $\frac{M_r(0)}{c_3 + M_r(0) + M_r(\Delta f)}$ | $\frac{M_r(\Delta f)}{c_3 + M_r(0) + M_r(\Delta f)}$ |
| | P+F+N | $M_w(\Delta p)$<br>$= c_1 \exp\left(-\frac{\Delta p}{c_2}\right)$ | $\frac{\hat{\sigma}_T}{M_w(0)}$ | $\frac{\hat{\sigma}_D}{M_w(\Delta p)}$ | $M_r(\Delta f)$<br>$= \exp\left(-\frac{\Delta f^2}{c_3}\right)$ | $\frac{M_r(0)}{c_4 + M_r(0) + M_r(\Delta f)}$ | $\frac{M_r(\Delta f)}{c_4 + M_r(0) + M_r(\Delta f)}$ |
| No normalization | F+P | $M_r(\Delta f)$<br>$= c_1 \exp\left(-\frac{\Delta f^2}{c_2}\right)$ | $\frac{\hat{\sigma}_T}{M_r(0)}$ | $\frac{\hat{\sigma}_D}{M_r(\Delta f)}$ | $M_w(\Delta p)$<br>$= c_3 \exp\left(-\frac{\Delta p}{c_4}\right)$ | $M_w(0)$ | $M_w(\Delta p)$ |
| | FP+FP | $M_r(\Delta f, \Delta p) =$<br>$c_1 \exp\left[-\left(\frac{\Delta f^2}{c_2} + \frac{\Delta p}{c_3}\right)\right]$ | $\frac{\hat{\sigma}_T}{M_r(0,0)}$ | $\frac{\hat{\sigma}_D}{M_r(\Delta f, \Delta p)}$ | $M_w(\Delta f, \Delta p) =$<br>$c_4 \exp\left[-\left(\frac{\Delta f^2}{c_5} + \frac{\Delta p}{c_6}\right)\right]$ | $M_w(0,0)$ | $M_w(\Delta f, \Delta p)$ |
| | FP+P | $M_r(\Delta f, \Delta p) =$<br>$c_1 \exp\left[-\left(\frac{\Delta f^2}{c_2} + \frac{\Delta p}{c_3}\right)\right]$ | $\frac{\hat{\sigma}_T}{M_r(0,0)}$ | $\frac{\hat{\sigma}_D}{M_r(\Delta f, \Delta p)}$ | $M_w(\Delta p)$<br>$= c_4 \exp\left(-\frac{\Delta p}{c_5}\right)$ | $M_w(0)$ | $M_w(\Delta p)$ |
| | FP:P | $M_r(\Delta f, \Delta p) =$<br>$c_1 \exp\left[-\left(\frac{\Delta f^2}{c_2} + \frac{\Delta p}{c_3}\right)\right]$ | $\frac{\hat{\sigma}_T}{M_r(0,0)}$ | $\frac{\hat{\sigma}_D}{M_r(\Delta f, \Delta p)}$ | $M_w(\Delta p)$<br>$= c_4 \exp\left(-\frac{\Delta p}{c_3}\right)$ | $M_w(0)$ | $M_w(\Delta p)$ |
| | FP:FP | $M_r(\Delta f, \Delta p) =$<br>$c_1 \exp\left[-\left(\frac{\Delta f^2}{c_2} + \frac{\Delta p}{c_3}\right)\right]$ | $\frac{\hat{\sigma}_T}{M_r(0,0)}$ | $\frac{\hat{\sigma}_D}{M_r(\Delta f, \Delta p)}$ | $M_w(\Delta f, \Delta p)$<br>$= M_r(\Delta f, \Delta p)$ | $M_w(0,0)$ | $M_w(\Delta f, \Delta p)$ |
| | P | $M_w(\Delta p)$<br>$= c_1 \exp\left(-\frac{\Delta p}{c_2}\right)$ | $\frac{\hat{\sigma}_T}{M_w(0)}$ | $\frac{\hat{\sigma}_D}{M_w(\Delta p)}$ | NA | $M_w(0)$ | $M_w(\Delta p)$ |
| | F | $M_r(\Delta f)$<br>$= c_1 \exp\left(-\frac{\Delta f^2}{c_2}\right)$ | $\frac{\hat{\sigma}_T}{M_r(0)}$ | $\frac{\hat{\sigma}_D}{M_r(\Delta f)}$ | NA | $M_r(0)$ | $M_r(\Delta f)$ |
| | P+F | $M_w(\Delta p)$<br>$= c_1 \exp\left(-\frac{\Delta p}{c_2}\right)$ | $\frac{\hat{\sigma}_T}{M_w(0)}$ | $\frac{\hat{\sigma}_D}{M_w(\Delta p)}$ | $M_r(\Delta f)$<br>$= c_3 \exp\left(-\frac{\Delta f^2}{c_4}\right)$ | $M_r(0)$ | $M_r(\Delta f)$ |
| Optimal | FP | $M_r(\Delta f, \Delta p) = c_1 \exp \left[ - \left( \frac{\Delta f^2}{c_2} + \frac{\Delta p}{c_3} \right) \right]$ | $\frac{\hat{\sigma}_T}{M_r(0,0)}$ | $\frac{\hat{\sigma}_D}{M_r(\Delta f, \Delta p)}$ | NA | $\frac{\sigma_D^2}{\sigma_T^2 + \sigma_D^2}$ | $\frac{\sigma_T^2}{\sigma_T^2 + \sigma_D^2}$ |
| | P | $M_w(\Delta p) = c_1 \exp \left( - \frac{\Delta p}{c_2} \right)$ | $\frac{\hat{\sigma}_T}{M_w(0)}$ | $\frac{\hat{\sigma}_D}{M_w(\Delta p)}$ | NA | $\frac{\sigma_D^2}{\sigma_T^2 + \sigma_D^2}$ | $\frac{\sigma_T^2}{\sigma_T^2 + \sigma_D^2}$ |
| | F | $M_r(\Delta f) = c_1 \exp \left( - \frac{\Delta f^2}{c_2} \right)$ | $\frac{\hat{\sigma}_T}{M_r(0)}$ | $\frac{\hat{\sigma}_D}{M_r(\Delta f)}$ | $M_r(\Delta f) = c_1 \exp \left( - \frac{\Delta f^2}{c_2} \right)$ | $\frac{\sigma_D^2}{\sigma_T^2 + \sigma_D^2}$ | $\frac{\sigma_T^2}{\sigma_T^2 + \sigma_D^2}$ |

$M_r$ : Reliability modulation function; $M_w$ : Weight modulation function; $\Delta f$ : Frequency disparity between target and distractor; $\Delta p$ : Position disparity between target and distractor; $\hat{\sigma}_T$ and $\hat{\sigma}_D$ : standard deviation of the representations of target and distractor stimuli, respectively, before modulation; $\sigma_T$ and $\sigma_D$ : standard deviation of the representations of target and distractor stimuli, respectively, after modulation; $W_T$ and $W_D$ : Weights to the target and distractor cue, respectively. $c$ indicates free parameter.
*Model abbreviations:*

**Supplemental Table 2.** Comparison of frequency combination models

| Models | | LL | LRT | AIC | BIC | Error in CP | Error in PSE | Error in JND | $\sqrt{(PSE^2 + JND^2)}$ | BF |
| --- | --- | --- | --- | --- | --- | --- | --- | --- | --- | --- |
| Normalization | F+P | -580.6±19.1 | 0 | 1169.1±38.1 | 1177.7±38.1 | 0.18±0.01 | 8.0±1.0 | 8.9±0.9 | 12.2±1.0 | 2.67 |
|  | FP+FP | -589.0±20.0 | -0.03±0.01 | 1190.0±40.0 | 1202.8±40.0 | 0.21±0.02 | 10.4±1.8 | 8.8±1.1 | 13.9±1.9 | 30.47 |
|  | FP+P | -586.1±20.2 | -0.02±0.01 | 1182.2±40.4 | 1192.9±40.4 | 0.20±0.01 | 9.8±1.1 | 9.7±1.0 | 14.0±1.2 | 20.58 |
|  | FP:P | -599.2±18.8 | -0.06±0.01 | 1206.4±37.7 | 1215.0±37.7 | 0.26±0.01 | 17.3±1.2 | 17.3±1.5 | 24.7±1.6 | 1728.8 |
|  | FP:FP | -583.1±18.6 | -0.01±0.01 | 1174.2±37.2 | 1182.8±37.2 | 0.19±0.01 | 7.4±0.9 | 10.4±1.3 | 13.0±1.3 | 7.84 |
|  | P | -593.0±19.3 | -0.04±0.01 | 1192.0±38.6 | 1198.4±38.6 | 0.22±0.01 | 10.6±1.0 | 14.3±0.9 | 18.0±0.9 | 174.19 |
|  | F | -584.8±18.3 | -0.02±0.01 | 1175.6±36.6 | 1182.1±36.6 | 0.20±0.01 | 9.9±1.1 | 9.7±0.8 | 14.0±1.1 | 11.43 |
|  | P+F | -588.0±17.6 | -0.03±0.01 | 1184.0±35.3 | 1192.6±35.3 | 0.21±0.01 | 9.7±1.1 | 12.4±1.1 | 15.9±1.3 | 72.9 |
| No normalization | F+P | -600.4±17.5 | -0.07±0.01 | 1208.8±35.0 | 1217.3±35.0 | 0.26±0.01 | 16.0±0.9 | 12.0±1.4 | 20.2±1.2 | 1734 |
|  | FP+FP | -618.6±18.5 | -0.13±0.01 | 1249.2±37.0 | 1262.0±37.0 | 0.31±0.01 | 19.0±1.2 | 20.9±3.8 | 28.9±3.2 | 1450.2 |
|  | FP+P | -638.6±25.5 | -0.19±0.04 | 1287.3±51.0 | 1298.0±51.0 | 0.36±0.04 | 25.1±3.4 | 33.3±8.8 | 42.5±8.9 | 36.02 |
|  | FP:P | -604.1±20.1 | -0.08±0.02 | 1216.1±40.2 | 1224.7±40.2 | 0.27±0.01 | 17.5±1.4 | 21.0±3.8 | 27.9±3.5 | 1107.9 |
|  | FP:FP | -599.2±17.6 | -0.06±0.01 | 1204.4±35.2 | 1210.9±35.2 | 0.25±0.01 | 14.0±1.3 | 10.4±1.0 | 17.9±0.7 | 3355.5 |
|  | P | -604.4±18.2 | -0.08±0.01 | 1212.8±36.3 | 1217.1±36.3 | 0.26±0.01 | 15.4±1.0 | 10.4±1.0 | 18.9±0.6 | 13896 |
|  | F | -602.8±17.8 | -0.08±0.01 | 1209.7±35.7 | 1214.0±35.7 | 0.26±0.01 | 15.5±1.5 | 10.4±1.0 | 19.2±0.8 | 12609 |
|  | P+F | -2063.9±167 | -2.44±0.17 | 4135.7±340 | 4144.3±340 | 0.29±0.01 | 18.5±1.5 | 15.6±0.7 | 24.5±1.1 | 9.2e16 |
| Optimal | FP | -587.8±18.1 | -0.03±0.01 | 1181.7±36.1 | 1188.1±36.1 | 0.21±0.01 | 10.8±1.2 | 10.4±1.1 | 15.2±1.3 | 55.52 |
|  | P | -588.2±18.0 | -0.03±0.01 | 1180.5±36.0 | 1184.8±36.0 | 0.21±0.01 | 10.2±1.1 | 10.7±1.1 | 15.0±1.3 | 19.16 |
|  | F | -588.7±18.2 | -0.03±0.01 | 1181.4±36.5 | 1185.7±36.5 | 0.21±0.01 | 11.2±1.1 | 10.0±1.2 | 15.1±1.5 | 58.91 |

Model comparison metrics estimated for models fit to each subject's data separately. Listed values indicate mean ± sem across subjects. Log-likelihood (LL), likelihood ratio test (LRT), Akaike information criteria (AIC), Bayesian information criteria (BIC), and prediction errors (SSE) in CP, PSE, and JND, and Bayes Factor (BF). The favored model under each criterion is indicated in the shaded cell.

**Supplemental Table 3.** Comparison of intensity combination models

| Models | LL | LRT | AIC | BIC | CP | PSE | BF |
| --- | --- | --- | --- | --- | --- | --- | --- |
| Normalization | -630.6±71.8 | 0 | 1263.1±143.5 | 1265.2±143.5 | 0.07±0.01 | 0.002±0.0003 | 0.43 |
| Cue averaging | -1140.2±89.2 | -1.2±0.2 | 2280.4±178.4 | 2280.4±178.4 | 0.52±0.05 | 0.014±0.000 | 782.87 |

Conventions as in Supplemental Table 2.

